# Feature Selection and Dimension Reduction for Single Cell RNA-Seq based on a Multinomial Model

**DOI:** 10.1101/574574

**Authors:** F. William Townes, Stephanie C. Hicks, Martin J. Aryee, Rafael A. Irizarry

## Abstract

Single cell RNA-Seq (scRNA-Seq) profiles gene expression of individual cells. Recent scRNA-Seq datasets have incorporated unique molecular identifiers (UMIs). Using negative controls, we show UMI counts follow multinomial sampling with no zero-inflation. Current normalization procedures such as log of counts per million and feature selection by highly variable genes produce false variability in dimension reduction. We propose simple multinomial methods, including generalized principal component analysis (GLM-PCA) for non-normal distributions, and feature selection using deviance. These methods outperform current practice in a downstream clustering assessment using ground-truth datasets.

## 1. Background

Single cell RNA-Seq (scRNA-Seq) is a powerful tool for profiling gene expression patterns in individual cells, facilitating a variety of analyses such as identification of novel cell types [1, 2]. In a typical protocol, single cells are isolated in liquid droplets and messenger RNA (mRNA) is captured from each cell, converted to cDNA by reverse transcriptase (RT), then amplified using polymerase chain reaction (PCR) [3, 4, 5]. Finally, fragments are sequenced and expression of a gene in a cell is quantified by the number of sequencing reads that mapped to that gene [6]. A crucial difference between scRNA-Seq and traditional bulk RNA-Seq is the low quantity of mRNA isolated from individual cells, which requires a larger number of PCR cycles to produce enough material for sequencing (bulk RNA-Seq comingles thousands of cells per sample). Thus, many of the reads counted in scRNA-Seq are duplicates of a single mRNA molecule in the original cell [7]. Early scRNA-Seq studies using protocols such as SMART-Seq2 [8] analyzed these *read counts* directly, and several methods were developed to facilitate this [9]. However, newer protocols typically include unique molecular identifiers (UMIs) which enable computational removal of PCR duplicates [10], producing *UMI counts*. Although a zero UMI count is equivalent to a zero read count, nonzero read counts are larger than their corresponding UMI counts. In general, all scRNA-Seq data contain large numbers of zero counts (often *>* 90% of the data, sometimes called *dropouts*). Here, we focus on the analysis of scRNA-Seq data with UMI counts.

Starting from raw counts, a scRNA-Seq data analysis typically includes normalization, feature selection, and dimension reduction steps. Normalization seeks to adjust for differences in experimental conditions between samples (individual cells), so that these do not confound true biological differences. For example, the efficiency of mRNA capture and RT is variable between samples (technical variation), causing different cells to have different total UMI counts, even if the number of molecules in the original cells is identical. Feature selection refers to excluding uninformative genes such as those which exhibit no meaningful biological variation across samples. Since scRNA-Seq experiments usually examine cells within a single tissue, only a small fraction of genes are expected to be informative since many genes are biologically variable only across different tissues. Dimension reduction aims to embed each cell’s high-dimensional expression profile into a low-dimensional representation to facilitate visualization and clustering.

While a plethora of methods [11, 12, 13, 5, 14] have been developed for each of these steps, here we describe what is considered to be the standard pipeline [14]. First, raw counts are normalized by scaling of sample-specific *size factors*, followed by log-transformation, which attempts to reduce skewness. Next, feature selection involves identifying the top 500-2,000 genes by computing either their coefficient of variation (highly variable genes [15, 16]), or average expression level (highly expressed genes) across all cells [14]. Alternatively, highly dropout genes may be retained [17]. Principal component analysis (PCA) [18] is the most popular dimension reduction method (see for example tutorials for Seurat [16] and Cell Ranger [5]). PCA compresses each cell’s 2,000-dimensional expression profile into, say, a 10-dimensional vector of principal component coordinates or latent factors. Prior to PCA, data are usually centered and scaled so that each gene has mean zero and standard deviation one (*z-score* transformation). Finally, a clustering algorithm can be applied to group cells with similar representations in the low-dimensional PCA space.

Despite the appealing simplicity of this standard pipeline, the characteristics of scRNA-Seq UMI counts present difficulties at each stage. Many normalization schemes derived from bulk RNA-Seq cannot compute size factors stably in the presence of large numbers of zeros [19]. A numerically stable and popular method is to set the size factor for each cell as the total counts divided by 106 (*counts per million*, CPM). Note that CPM does not alter zeros, which dominate scRNA-Seq data. Log-transformation is not possible for exact zeros so it is common practice to add a small *pseudocount* such as one to all normalized counts prior to taking the log. The choice of pseudocount is arbitrary and can introduce subtle biases in the transformed data [20]. For a statistical interpretation of the pseudocount see Section 4.2. Similarly, the use of highly variable genes for feature selection is somewhat arbitrary since the observed variability will depend on the pseudocount: pseudocounts close to zero arbitrarily increase the variance of genes with zero counts. Finally, PCA implicitly relies on Euclidean geometry, which may not be appropriate for highly sparse, discrete, and skewed data, even after normalizations and transformations [21].

Widely used methods for analysis of scRNA-Seq lack statistically rigorous justification based on a plausible data generating mechanism for UMI counts. Instead, it appears many of the techniques have been borrowed from the data analysis pipelines developed for read counts, especially those based on bulk RNA-Seq [22]. For example, models based on the lognormal distribution cannot account for exact zeros, motivating the development of zero-inflated lognormal models for scRNA-Seq read counts [23, 24, 25, 26]. Alternatively, ZINB-WAVE uses a zero-inflated negative binomial model for dimension reduction of read counts [27]. However, as shown below, the sampling distribution of UMI counts differs markedly from read counts, so application of read count models to UMI counts needs either theoretical or empirical justification.

We present a unifying statistical foundation for scRNA-Seq with UMI counts based on the multinomial distribution. The multinomial model adequately describes negative control data and there is no need to model zero inflation. We show the mechanism by which PCA on log-normalized UMI counts can lead to distorted low-dimensional factors and false discoveries. We identify the source of the frequently observed and undesirable fact that the fraction of zeros reported in each cell drives the first principal component in most experiments [28]. To remove these distortions, we propose the use of GLM-PCA, a generalization of PCA to exponential family likelihoods [29]. GLM-PCA operates on raw counts, avoiding the pitfalls of normalization. We also demonstrate that applying PCA to deviance or Pearson residuals provides a useful and fast approximation to GLM-PCA. We provide a closed-form deviance statistic as a feature selection method. We systematically compare the performance of all combinations of methods using three ground-truth datasets and assessment procedures from [14]. We conclude by suggesting best practices.

## 2. Results and discussion

### 2.1 Datasets

We used five public 10x genomics UMI counts datasets from [5] to benchmark our methods. The first dataset is a highly controlled experiment specifically designed to understand technical variability. No actual cells were used to generate this dataset. Instead, each of the 1,015 droplets received the same ratio of 92 synthetic spike-in RNA molecules from External RNA Controls Consortium (ERCC). We refer to this dataset as the *technical replicates negative control* as there is no biological variability whatsoever and, in principle, each expression profile should be the same.

The second dataset was generated by processing a homogeneous population of 2,612 monocyte cells. The cells were purified using fluorescence activated cell sorting (FACS). We refer to this dataset as the *biological replicates negative control*. Because these cells were all the same type, we did not expect to observe any significant differences in unsupervised analysis.

The third dataset consists of 68,579 fresh, unsorted peripheral blood mononuclear cells (PBMCs). This dataset was used to compare computational speed of different dimension reduction algorithms. We refer to it as the PBMC 68K dataset.

The remaining two datasets were created by [14]. In the Zheng 4eq dataset, there are 3,994 PBMCs divided equally into four cell types. In the Zheng 8eq dataset, there are 3,994 PBMCs divided equally into eight cell types. In these positive control datasets, the cluster identity of all cells was assigned independently of gene expression (using FACS), so they served as ground truth labels.

### 2.2 UMI count distribution differs from reads

To illustrate the marked difference between UMI count distributions and read count distributions, we created histograms from individual genes and spike-ins of the negative control data. Here, the UMI counts are the computationally deduplicated versions of the read counts; both measurements are from the same experiment, so no differences are due to technical or biological variation. The results suggest that while read counts appear zero-inflated and multimodal, UMI counts follow a discrete distribution with no zero inflation (Figure S1). The apparent zero inflation in read counts is a result of PCR duplicates.

### 2.3 Multinomial sampling distribution for UMI counts

Consider a single cell *i* containing *t*_*i*_ total mRNA transcripts. Let *n*_*i*_ be the total number of UMIs for the same cell. When the cell is processed by a scRNASeq protocol, it is lysed, then some fraction of the transcripts are captured by beads within the droplets. A series of complex biochemical reactions occur, including attachment of barcodes and UMIs, and reverse transcription of the captured mRNA to a cDNA molecule. Finally, the cDNA is sequenced and PCR duplicates are removed to generate the UMI counts [5]. In each of these stages, some fraction of the molecules from the previous stage are lost [30, 5, 7]. In particular, reverse transcriptase is an inefficient and error-prone enzyme [31]. Therefore the number of UMI counts representing the cell is much less than the number of transcripts in the original cell (*n*_*i*_ *≪ t*_*i*_). Specifically, *n*_*i*_ typically ranges from 1, 000 −10, 000 while *t*_*i*_ is estimated to be approximately 200, 000 for a typical mammalian cell [32]. Furthermore, which molecules are selected and successfully become UMIs is a random process. Let *x*_*ij*_ be the true number of mRNA transcripts of gene *j* in cell *i*, and *y*_*ij*_ be the UMI count for the same gene and cell. We define the *relative abundance π*_*ij*_ as the true number of mRNA transcripts represented by gene *j* in cell *i* divided by the total number of mRNA transcripts in cell *i*. Relative abundance is given by *π*_*ij*_ = *x*_*ij*_ */t*_*i*_ where total transcripts *t*_*i*_ = Σ_*j*_ *x*_*ij*_. Since *n*_*i*_ *≪ t*_*i*_, there is a “competition to be counted” [33]; genes with large relative abundance *π*_*ij*_ in the original cell are more likely to have nonzero UMI counts, but genes with small relative abundances may be observed with UMI counts of exact zeros. The UMI counts *y*_*ij*_ are a multinomial sample of the true biological counts *x*_*ij*_, containing only relative information about expression patterns in the cell [34, 33].

The multinomial distribution can be approximated by independent Poisson distributions, and overdispersed multinomials by independent negative binomial distributions. These approximations are useful for computational tractability. Details are provided in the Methods.

The multinomial model makes two predictions which we verified using negative control data. First, the fraction of zeros in a sample (cell or droplet) is inversely related to the total number of UMIs in that sample. Second, the probability of an endogenous gene or ERCC spike-in having zero counts is a decreasing function of its mean expression (equations provided in Methods). Both of these predictions were validated by the negative control data (Figure 1). In particular, the empirical probability of a gene being zero across droplets was well calibrated to the theoretical prediction based on the multinomial model. This also demonstrates that UMI counts are not zero inflated.

**Figure 1:**
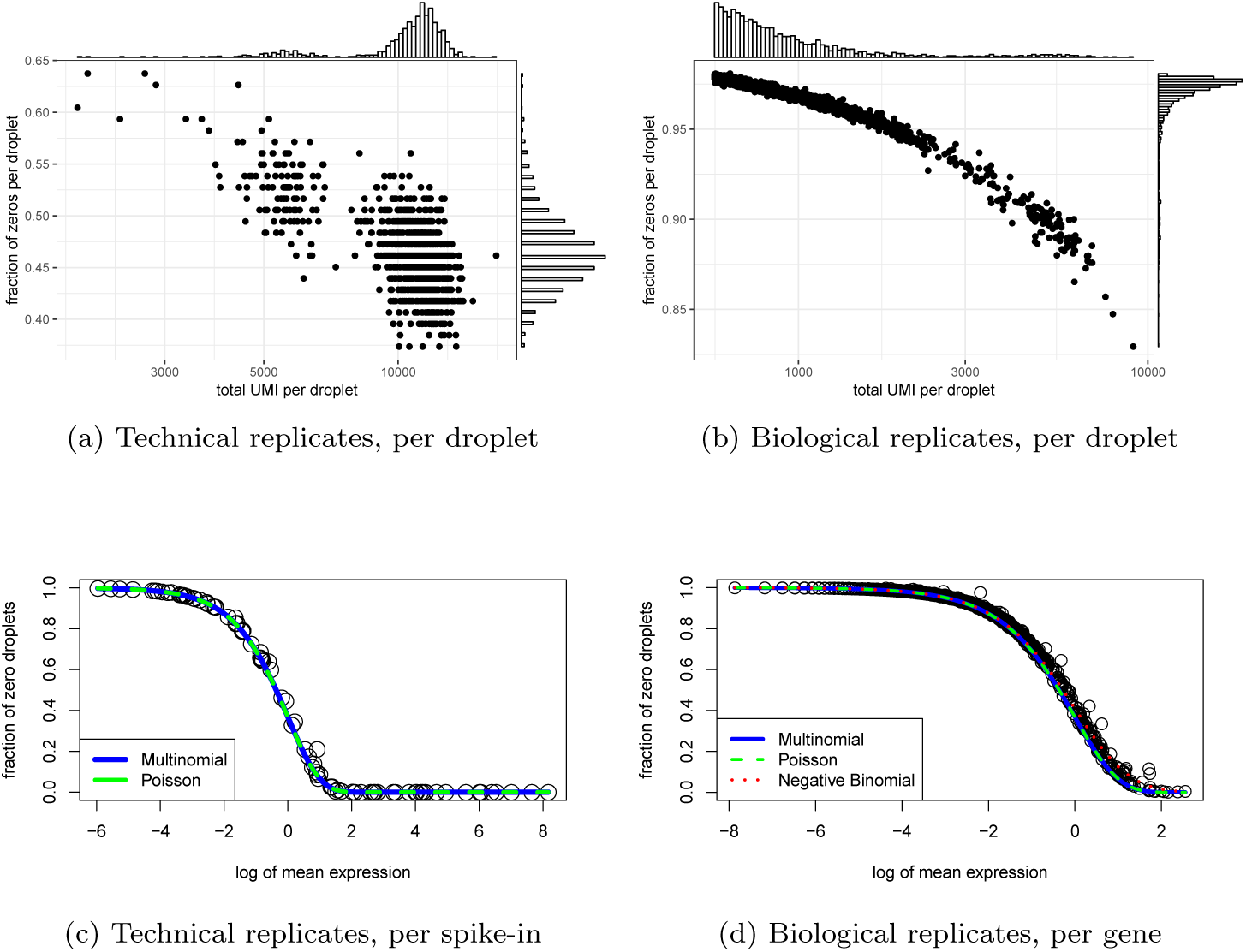
Multinomial model adequately characterizes sampling distributions of technical and biological replicates negative control data. a) Fraction of zeros is plotted against the total number of UMI in each droplet for the technical replicates. b) As a) but for cells in the biological replicates. c) After down-sampling replicates to 10,000 UMIs per droplet to remove variability due to differences in sequencing depth, the fraction of zeros is computed for each gene and plotted against the log of expression across all samples for the technical replicates data. The solid curve is theoretical probability of observing a zero as a function of the expected counts derived from the multinomial model (blue) and its Poisson approximation (green). d) As c) but for the biological replicates dataset and after down-sampling to 575 UMIs per cell. Here we also add the theoretical probability derived from a negative binomial model (red).

These results are consistent with [35], which also found that the relationship between average expression and zero probability follows the theoretical curve predicted by a Poisson model using negative control data processed with Indrop [4] and Dropseq [3] protocols. This suggests our data generating mechanism is an accurate model of technical noise in real data.

### 2.4 Normalization and log transformation distorts UMI data

Standard scRNA-Seq analysis involves normalizing raw counts using size factors, applying a log transformation with a pseudocount, and then centering and scaling each gene before dimension reduction. The most popular normalization is counts per million (CPM). The CPM are defined as (*y*_*ij*_ */n*_*i*_)*10^6^ (i.e. the size factor is *n*_*i*_ */*10^6^). This is equivalent to the MLE for relative abundance 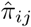 multiplied by 10^6^. The log-CPM are then 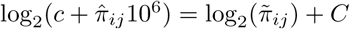, where 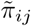 is a maximum a posteriori estimator (MAP) for *π*_*ij*_ (mathematical justification and interpretation of this approach provided in Methods). The additive constant *C* is irrelevant if data are centered for each gene after log transformation, as is common practice. Thus, normalization of raw counts is equivalent to using MLEs or MAP estimators of the relative abundances.

Log transformation of MLEs is not possible for UMI counts due to exact zeros, while log transformation of MAP estimators of *π*_*ij*_ systematically distorts differences between zero and nonzero UMI counts, depending on the arbitrary pseudocount *c* (derivations provided in Methods). To illustrate this phenomenon, we examined the distribution of an illustrative gene before and after the log transform with varying normalizations using the biological replicates negative control data (Figure 2). Consistent with our theoretical predictions, this artificially caused the distribution to appear zero inflated and exaggerated differences between cells based on whether the count was zero or nonzero.

**Figure 2:**
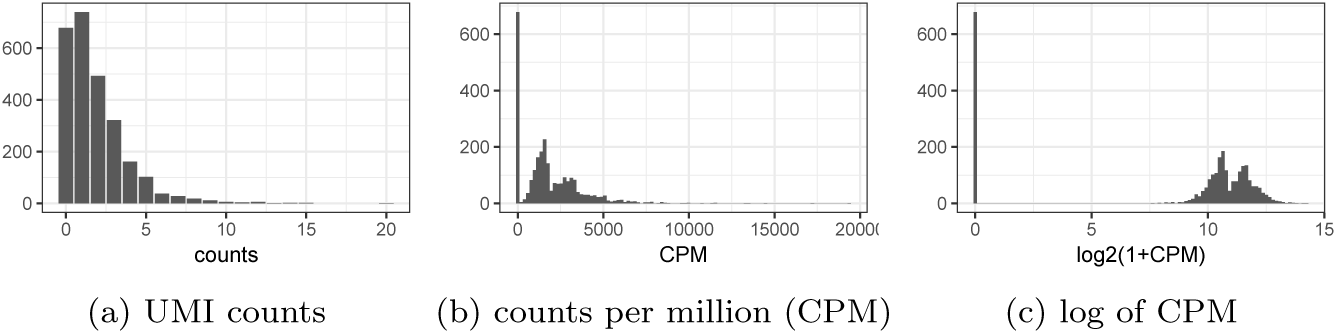
Example of how current approaches to normalization and transformation artificially distort differences between zero and nonzero counts. a) UMI count distribution for gene ENSG00000114391 in the biological replicates negative control dataset. b) Counts per million (CPM) distribution for the exact same count data. c) Distribution of *log*_2_(1 + *CPM*) values for the exact same count data.

Focusing on the entire negative control datasets, we applied PCA to the log transformed CPMs and observed a strong correlation between the first principal component (PC) and the fraction of zeros, consistent with [28]. Additionally, the first PC correlates with the log of total UMI, which is consistent with the multinomial model (Figure 3). Based on these results, the log transformation is not necessary and in fact detrimental for analysis of UMI counts. The benefits of avoiding normalization by instead directly modeling raw counts have been demonstrated in the context of differential expression [36]. Where normalization is unavoidable, we propose the use of approximate multinomial deviance residuals (defined in Section 4.4) instead of log-transformed CPM.

**Figure 3:**
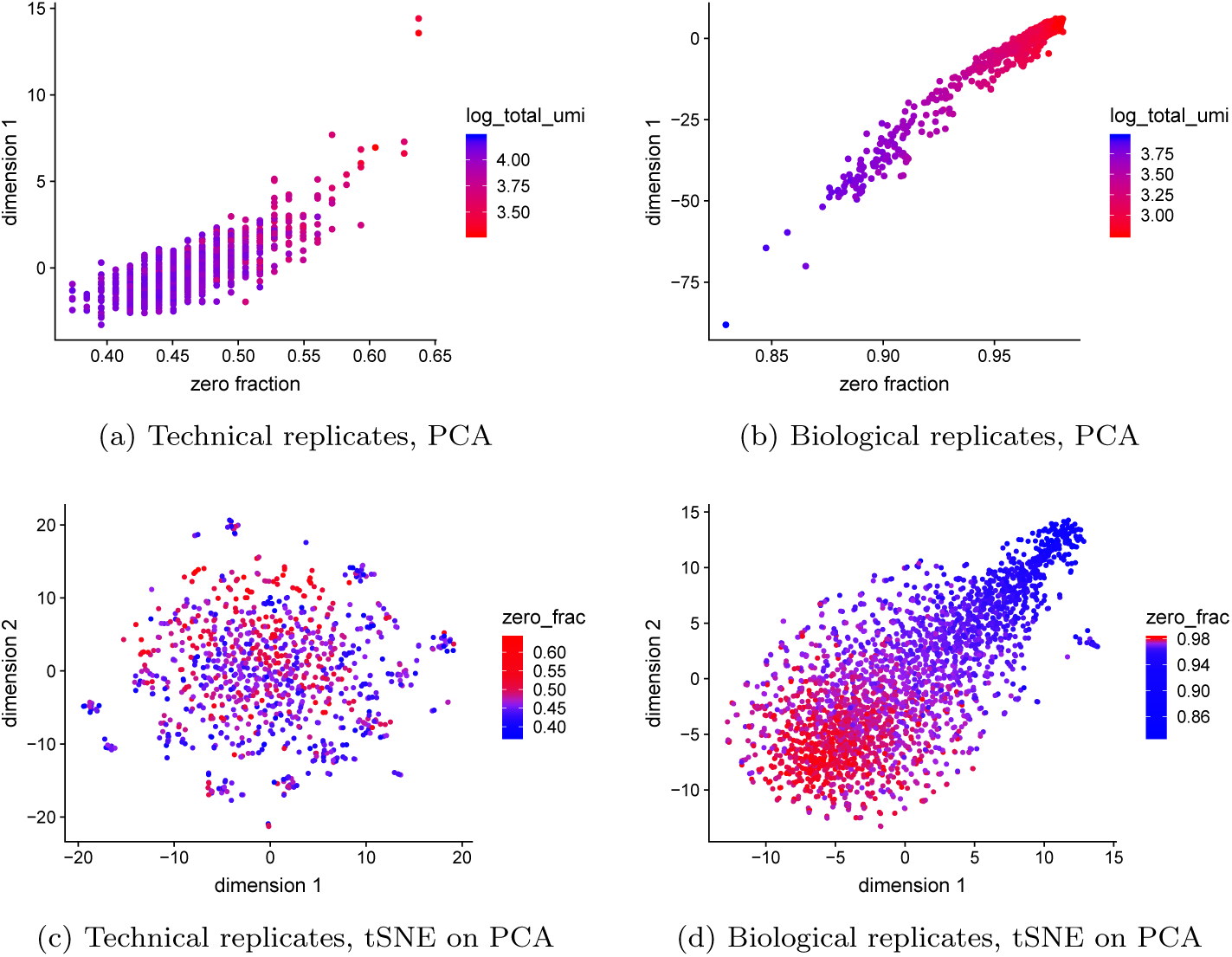
Current approaches to normalization and transformation induce variability in the fraction of zeros across cells to become the largest source of variability which in turn biases clustering algorithms to produce false positive results based on distorted latent factors. a) First principal component (PC) from the technical replicates dataset plotted against fraction of zeros for each cell. A red to blue color scale represents total UMIs per cell. b) as a) but for the biological replicates data. c) Using the technical replicates, we applied t-distributed stochastic neighbor embedding (tSNE) with perplexity 30 to the top 50 PCs computed from log-CPM. The first two tSNE dimensions are shown with a blue to red color scale representing the fraction of zeros. d) as c) but for the biological replicates data. Here we do not expect to find differences, yet we see distorted latent factors being driven by the total UMIs. PCA was applied to 5,000 random genes.

### 2.5 Zero inflation is an artifact of log-normalization

To see how normalization and log-transformation introduce the appearance of zero inflation, consider the following example. Let *y*_*ij*_ be the observed UMI counts following a multinomial distribution with size *n*_*i*_ for each cell and relative abundance *π*_*j*_ for each gene, constant across cells. Focusing on a single gene *j, y*_*ij*_ follows a binomial distribution with parameters *n*_*i*_, *p*_*j*_. Assume *π*_*j*_ = 10^−4^ and the *n*_*i*_ range from 1, 000 −3, 000, which is consistent with the biological replicates negative control data (Figures S1 and 1). Under this assumption we expect to see about 74-90% zeros, 22-30% ones, and less than 4% values above one. However, notice that after normalization to CPM and log transformation, all the zeros remain log 2(1 + 0) = 0, yet the ones turn into values ranging from log^2^(1+ 1*/*3000 *\** 10^6^) = log_2_(334) ≈ 8.4 to log^2^(1001) ≈ 10. The few values that are 2 will have values ranging from log^2^(668) ≈9.4 to log_2_(2001) ≈11. The large, artificial gap between zero and nonzero values makes the log-normalized data appear zero-inflated (Figure 2). The variability in CPM values across cells is almost completely driven by the variability in *n*_*i*_. Indeed, it shows up as the primary source of variation in PCA plots (Figure 3).

### 2.6 Generalized PCA for dimension reduction of sparse counts

While PCA is a popular dimension reduction method, it is implicitly based on Euclidean distance, which corresponds to maximizing a Gaussian likelihood. Since UMI counts are not normally distributed, even when normalized and log transformed, this distance metric is inappropriate, causing PCA to produce distorted latent factors (Figure 3). We propose the use of PCA for generalized linear models (GLMs) [29], or GLM-PCA as a more appropriate alternative. The GLM-PCA framework allows for a wide variety of likelihoods suitable for data types such as counts and binary values. While the multinomial likelihood is ideal for modeling technical variability in scRNA-Seq UMI counts (Figure 1), in many cases there may be excess biological variability present as well. For example, if we wish to capture variability due to clusters of different cell types in a dimension reduction, we may wish to exclude biological variability due to cell cycle. Biological variability not accounted for by the sampling distribution may be accomodated by using a Dirichlet-multinomial likelihood, which is overdispersed relative to the multinomial. In practice, both the multinomial and Dirichlet-multinomial are computationally intractable, and may be approximated by the Poisson and negative binomial likelihoods, respectively (detailed derivations provided in Methods). We found that numerical convergence of GLM-PCA with negative binomial likelihood was unstable, due to the difficulty of estimating the dispersion parameter, so we focused on the simpler Poisson likelihood in our assessments. Intuitively, using Poisson instead of negative binomial implies we assume the biological variability is captured by the factor model and the unwanted biological variability is small relative to the sampling variability. We ran Poisson GLM-PCA on the technical and biological replicates negative control datasets and found it removed the spurious correlation between the first dimension and the total UMIs and fraction of zeros (Figure 4).

**Figure 4:**
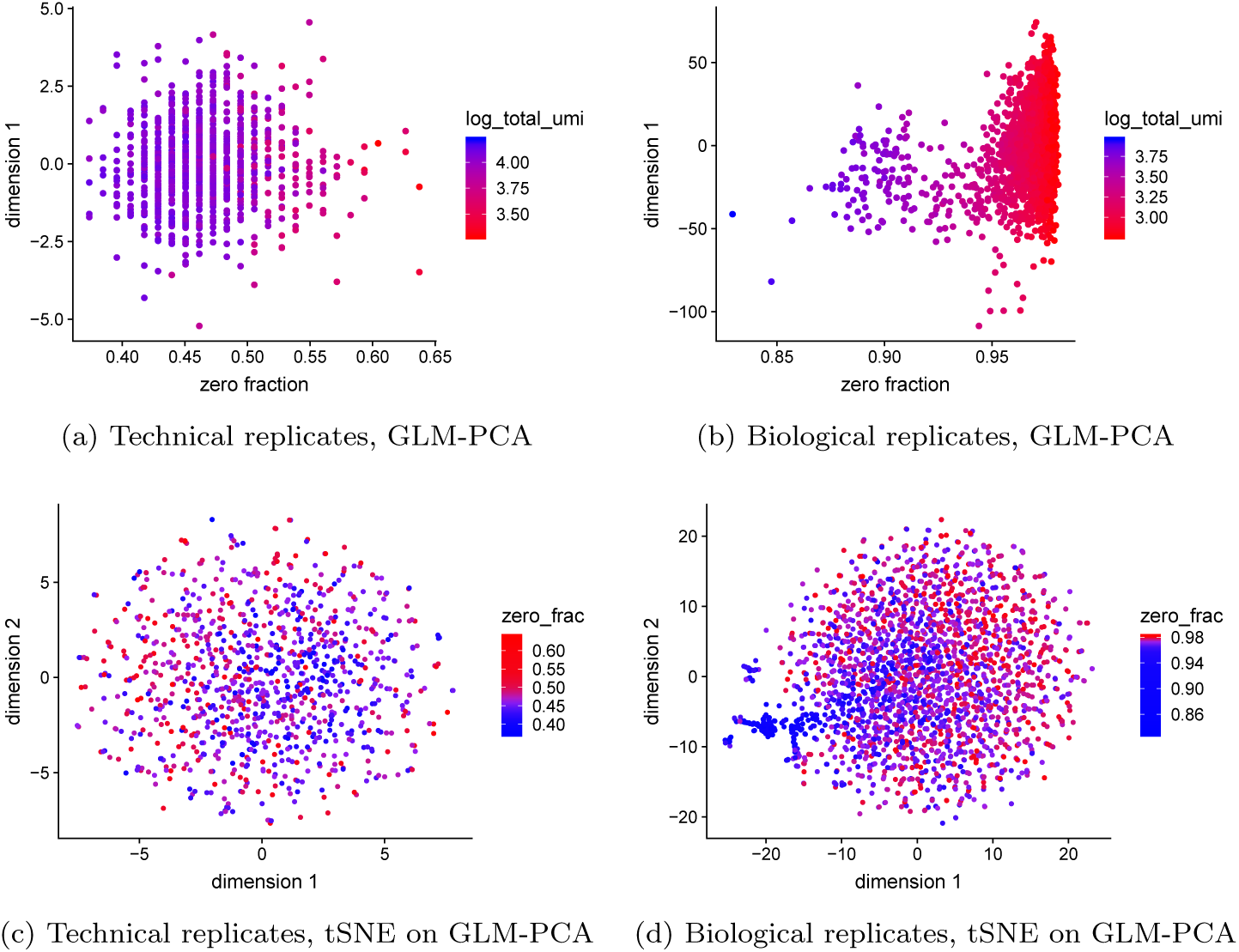
GLM-PCA dimension reduction is not affected by unwanted fraction of zeros variability and avoids false positive results. a) First GLM-PCA dimension (analogous to first principal component) plotted against the fraction of zeros for the technical replicates with colors representing the total UMIs. b) as a) but using biological replicates. c) Using the technical replicates, we applied t-distributed stochastic neighbor embedding (tSNE) with perplexity 30 to the top 50 GLM-PCA dimensions. The first two tSNE dimensions are shown with a blue to red color scale representing the fraction of zeros. d) as c) but for the biological replicates data. GLM-PCA using the Poisson approximation to the multinomial was applied to the same 5,000 random genes as in Figure 3.

### 2.7 Deviance residuals provide fast approximation to GLMPCA

One disadvantage of GLM-PCA is it depends on an iterative algorithm to obtain estimates for the latent factors, and is at least ten times slower than PCA. We therefore propose a fast approximation to GLM-PCA. When using PCA a common first step is to center and scale the data for each gene as z-scores. This is equivalent to the following procedure. First, specify a null model of constant gene expression across cells, assuming a normal distribution. Next, find the MLEs of its parameters for each gene (the mean and variance). Finally, compute residuals of the model as the z-scores (derivation provided in Methods). The fact that scRNA-Seq data are skewed, discrete, and possessing many zeros suggests the normality assumption may be inappropriate. Further, using z-scores does not account for variability in total UMIs across cells. Instead, we propose to replace the normal null model with a multinomial null model as a better match to the data generating mechanism. The analogs to z-scores under this model are called deviance and Pearson residuals. Mathematical formulae are presented in Methods. Use of multinomial residuals enables a fast transformation similar to z-scores that avoids difficulties of normalization and log-transformation by directly modeling counts. Additionally, this framework allows straightforward adjustment for covariates such as cell cycle signatures or batch labels.

### 2.8 Feature selection using deviance

Feature selection, or identification of informative genes, may be accomplished by ranking genes using the *deviance*, which quantifies how well each gene fits a null model of constant expression across cells. Unlike the competing highly variable or highly expressed genes methods, which are sensitive to normalization, ranking genes by deviance operates on raw UMI counts. An approximate multinomial deviance statistic can be computed in closed-form (formula provided in the Methods). We compared gene ranks for all three feature selection methods (deviance, highly expressed, and highly variable genes) on the 8eq dataset, which contained eight different known cell types. We found strong concordance between highly deviant genes and highly expressed genes (Spearman’s rank correlation *r* = 0.9987), while highly variable genes correlated weakly with both high expression (*r* = 0.3835) and deviance (*r* = 0.3738); see also Figure S2.

### 2.9 Multinomial models improve unsupervised clustering

Dimension reduction with GLM-PCA or its fast multinomial residuals approximation improved clustering performance over competing methods (Figure 5a). Feature selection by multinomial deviance was superior to highly variable genes (Figure 5b).

**Figure 5:**
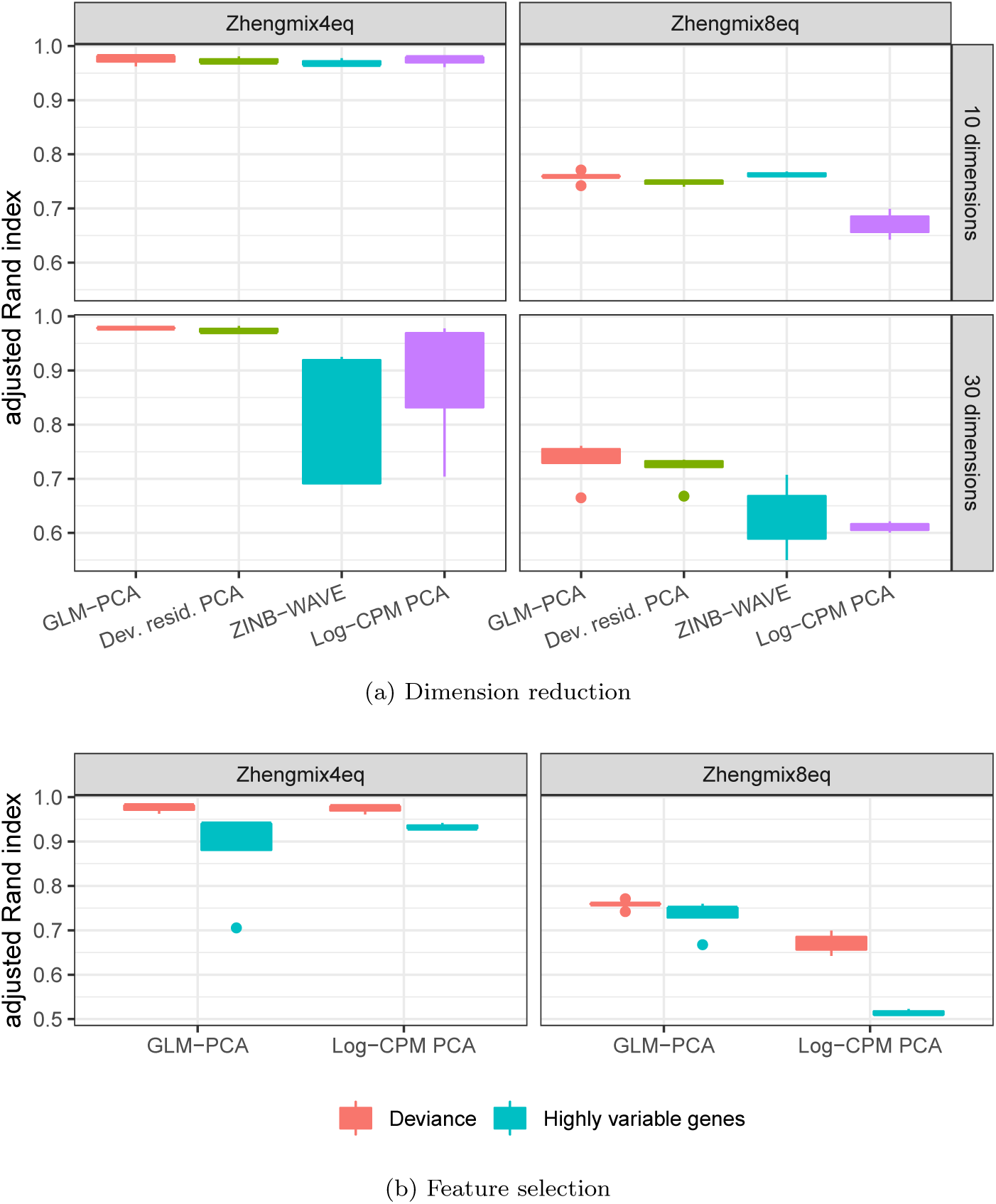
Dimension reduction with GLM-PCA and feature selection using deviance improves Seurat clustering performance. Each column represents a different groundtruth dataset from [14]. a) Comparison of dimension reduction methods based on the top 1,500 informative genes identified by approximate multinomial deviance. The Poisson approximation to the multinomial was used for GLM-PCA. Dev. resid. PCA: PCA on approximate multinomial deviance residuals. b) Comparison of feature selection methods. The top 1,500 genes identified by deviance and highly variable genes were passed to two different dimension reduction methods: GLM-PCA and PCA on log transformed CPM. Only results with the number of clusters within 25% of the true number are presented.

Using the two ground-truth datasets described in Section 2.1, we systematically compared the clustering performance of all combinations of previously described methods for normalization, feature selection, and dimension reduction. In addition, we compared against ZINB-WAVE since it also avoids requiring the user to pre-process and normalize the UMI count data (e.g. log transformation of CPM) and accounts for varying total UMIs across cells [27]. After obtaining latent factors, we used Seurat and k-means to infer clusters, and compared these to the known cell identities using Adjusted Rand Index (ARI, [37]). We varied the number of latent dimensions and number of clusters to assess robustness. Where possible, we used the same combinations of hyperparameters as [14] to facilitate comparisons to their extensive benchmarking (details are provided in Methods Section 4.6).

We compared the Seurat clustering performance of GLM-PCA (with Poisson approximation to multinomial) to running PCA on deviance residuals, which adhere more closely to the normal distribution than log-CPM. We found both of these approximate multinomial methods gave similar results on the 4eq dataset, and outperformed PCA on log-CPM z-scores. However, GLM-PCA outperformed the residuals method on the 8eq dataset. Also, performance on ZINB-WAVE factors degraded when the number of latent dimensions increased from 10 to 30, whereas GLM-PCA and its fast approximation with deviance residuals was robust to this change (Figure 5a). The performance of Pearson residuals was similar to that of deviance residuals (Figure S3).

Focusing on feature selection methods, deviance outperformed highly variable genes across both datasets and across dimension reduction methods (Figure 5b). Filtering by highly expressed genes led to similar clustering performance as deviance (Figure S3), because both criteria identified strongly overlapping gene lists for these data (Figure S2). The combination of feature selection with deviance and dimension reduction with GLM-PCA also improved clustering performance when k-means was used in place of Seurat (Figure S4). A complete table of results is publicly available (Section 5).

### 2.10 Computational efficiency of multinomial models

We measured time to convergence for reduction to two latent dimensions of GLM-PCA, ZINB-WAVE, PCA on log-CPM, PCA on deviance residuals, and PCA on Pearson residuals. Using the top 600 highly deviant genes, we subsampled the PBMC 68K dataset to 680, 6,800, and 68,000 cells. All methods scaled approximately linearly with increasing numbers of cells, but GLM-PCA was 23-63 times faster than ZINB-WAVE across sample sizes (Figure S5). Specifically, GLM-PCA processed 68,000 cells in less than seven minutes. The deviance and Pearson residuals methods exhibited speeds comparable to PCA: 9-26 times faster than GLM-PCA. We also timed dimension reduction of the 8eq dataset (3,994 cells) from 1,500 highly deviant genes to ten latent dimensions. PCA (with either log-CPM, deviance, or Pearson residuals) took 7 sec, GLM-PCA took 4.7 min, and ZINB-WAVE took 86.6 min.

## 3. Conclusions

We have outlined a statistical framework for analysis of scRNA-Seq data with UMI counts based on a multinomial model, providing effective and simple to compute methods for feature selection and dimension reduction. We found that UMI count distributions differ dramatically from read counts, are well-described by a multinomial distribution and are not zero-inflated. Log transformation of normalized UMI counts is detrimental, because it artificially exaggerates differences between zeros and all other values. For feature selection, or identification of informative genes, deviance is a more effective criterion than highly variable genes. Dimension reduction via GLM-PCA, or its fast approximation using residuals from a multinomial model, leads to better clustering performance than PCA on z-scores of log-CPM.

Although our methods were inspired by scRNA-Seq UMI counts, they may be useful for a wider array of data sources. Any high dimensional, sparse dataset where samples contain only relative information in the form of counts may conceivably be modeled by the multinomial distribution. Under such scenarios our methods are likely to be more effective than applying log-transformations and standard PCA. A possible example is microbiome data.

We have not addressed major topics in the scRNA-Seq literature such as pseudotime inference [38], differential expression [39], and spatial analysis [40]. However, the statistical ideas outlined here can also be used to improve methods in these more specialized types of analyses. In addition, adapting the GLM-PCA model to incorporate covariates such as batch labels or cell cycle signatures would be straightforward.

Our results have focused on (generalized) linear models for simplicity of exposition. Recently, several promising nonlinear dimension reductions for scRNA-Seq have been proposed. The variational autoencoder (VAE, a type of neural network) method scVI [41] utilizes a negative binomial likelihood in the decoder, while the encoder relies on log-normalized input data for numerical stability. The Gaussian process method tGPLVM [42] models log-transformed counts. In both cases, we suggest replacing log-transformed values with deviance residuals to improve performance. Nonlinear dimension reduction methods may also depend on feature selection to reduce memory consumption and speed computation; here, our deviance method may be utilized as an alternative to high variability for screening informative genes.

The statistical approaches described here have not been validated against scRNA-Seq data without UMIs, such as SMART-Seq2 and other plate protocols [8], since non-UMI data contain PCR duplicates. To apply the ideas to these data, one would need to be able to infer the UMI counts for data with PCR replicates [7].

## 4 Methods

### 4.1 Multinomial Model for scRNA-Seq

Let *y*_*ij*_ be the observed UMI counts for cell or droplet *i* and gene or spike-in *j*. Let *n*_*i*_ = Σ_*j*_ *y*_*ij*_ be the total UMIs in the sample, and *π*_*ij*_ be the unknown true relative abundance of gene *j* in cell *i*. The random vector **y**_*i*_ = (*y*_*i1*_, *…*, *y*_*iJ*_)^*T*^ with constraint Σ*_j_ y*_*ij*_ = *n*_*i*_ follows a multinomial distribution with density function

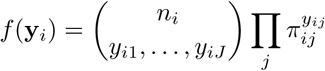

Focusing on a single gene *j* at a time, the marginal distribution of *y*_*ij*_ is binomial with parameters *n*_*i*_ and *π*_*ij*_. The marginal mean is E[*y*_*ij*_] = *n*_*i*_*π*_*ij*_ = *µ*_*ij*_, the marginal variance is var 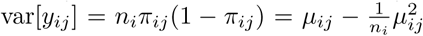, and the marginal probability of a zero count is 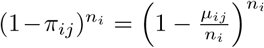. The correlation between two genes *j, k* is

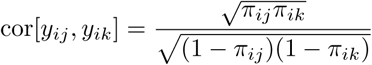

The correlation is induced by the sum to *n*_*i*_ constraint. As an extreme example, if there are only two genes (*J* = 2), increasing the count of the first gene automatically reduces the count of the second gene since they must add up to *n*_*i*_ under multinomial sampling. This means when *J* = 2 there is perfect anti-correlation between the gene counts which has nothing to do with biology. More generally, when either *J* or *n*_*i*_ is small, gene counts will be negatively correlated independent of biological gene-gene correlations, and it is not possible to analyze the data on a gene-by-gene basis (for example, by ranking and filtering genes for feature selection). Rather, comparisons are only possible between pairwise ratios of gene expression values [43]. Yet this type of analysis is difficult to interpret and computationally expensive for large numbers of genes (i.e. in high dimensions). Fortunately, under certain assumptions, more tractable approximations may be substituted for the true multinomial distribution.

First, note that if correlation is ignored, the multinomial may be approximated by *J* independent binomial distributions. Intuitively, this approximation will be reasonable if all *π*_*ij*_ are very small, which is likely to be satisfied for scRNA-Seq if the number of genes *J* is large, and no single gene constitutes the majority of mRNAs in the cell. If *n*_*i*_ is large and *π*_*ij*_ small, each binomial distribution can be further approximated by a Poisson with mean *n*_*i*_*π*_*ij*_. Alternatively, the multinomial can be constructed by drawing *J* independent Poisson random variables and conditioning on their sum. If *J* and *n*_*i*_ are large, the difference between the conditional, multinomial distribution and the independent Poissons becomes negligible. Since in practice *n*_*i*_ is large, the Poisson approximation to the multinomial may be reasonable [44, 45, 46, 47].

The multinomial model does not account for biological variability. As a result, an overdispersed version of the multinomial model may be necessary. This can be accommodated with the Dirichlet-multinomial distribution. Let **y**_*i*_ be distributed as a multinomial conditional on the relative abundance parameter vector *π*_*i*_ = (*π*_*i1*_, *…*, *π*_*iJ*_)^*T*^. If *π*_*i*_ is itself a random variable with symmetric Dirichlet distribution having shape parameter *α*, the marginal distribution of **y**_*i*_ is Dirichlet-multinomial. This distribution can itself be approximated by independent negative binomials. First, note that a symmetric Dirichlet random vector can be constructed by drawing *J* independent gamma variates with shape parameter *α* and dividing by their sum. Suppose (as above) we approximate the conditional multinomial distribution of **y**_*i*_ such that *y*_*ij*_ follows an approximate Poisson distribution with mean *n*_*i*_*π*_*ij*_. Let *λ*_*ij*_ be a collection of non-negative random variables such that 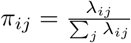 We require that *π*_*i*_ follow a symmetric Dirichlet, which is accomplished by having *λ*_*ij*_ follow indendent Gamma distributions with shape *α* and mean *n*_*i*_*/J*. This implies Σ_*j*_ *λ*_*ij*_ follows a Gamma with shape *Jα* and mean *n*_*i*_. As *J → ∞* this distributions converges to a point mass at *n*_*i*_, so for large *J* (satisfied by scRNA-Seq), Σ _*j*_ *λ*_*ij*_ *≈ n*_*i*_. This implies that *y*_*ij*_ approximately follows a conditional Poisson distribution with mean *λ*_*ij*_, where *λ*_*ij*_ is itself a Gamma random variable with mean *n*_*i*_*/J* and shape *α*. If we then integrate out *λ*_*ij*_ we obtain the marginal distribution of *y*_*ij*_ as negative binomial with shape *α* and mean *n*_*i*_*/J*. Hence a negative binomial model for count data may be regarded as an approximation to an overdispersed Dirichlet-multinomial model.

Parameter estimation with multinomial models (and their binomial or Poisson approximations) is straightforward. First, suppose we observe replicate samples **y**_*i*_, *i* = 1, *…*, *I* from the same underlying population of molecules, where the relative abundance of gene *j* is *π*_*j*_. This is a null model because it assumes each gene has a constant expected expression level and there is no biological variation across samples. Regardless of whether one assumes a multinomial, binomial, or Poisson model, the maximum likelihood estimator (MLE) of *π*_*j*_ is 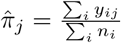 where *n*_*i*_ is the total count of sample *i*. In the more realistic case that relative abundances *π*_*ij*_ of genes vary across samples, the MLE is 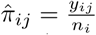

An alternative to the MLE is the maximum a posteriori (MAP) estimator. Suppose a symmetric Dirichlet prior with concentration parameter *α*_*i*_ is combined with the multinomial likelihood for cell *i*. The MAP estimator for *π*_*ij*_ is given by

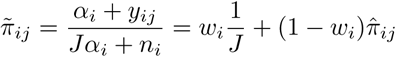

where *w*_*i*_ = *Jα*_*i*_*/*(*Jα*_*i*_ + *n*_*i*_), showing that the MAP is a weighted average of the prior mean that all genes are equally expressed (1*/J*) and the 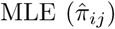. Compared to the MLE, the MAP biases the estimate toward the prior where all genes have the same expression. Larger values of *α*_*i*_ introduce more bias, while *α*_*i*_ *→* 0 leads to the MLE. If *α*_*i*_ *>* 0, the smallest possible value of 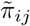 is *α*_*i*_*/*(*Jα*_*i*_ + *n*_*i*_) rather than zero for the MLE. When there are many zeros in the data, MAP can stabilize relative abundance estimates at the cost of introducing bias.

### 4.2 Mathematics of distortion from log-normalizing UMIs

Suppose the true counts in cell *i* are given by *x*_*ij*_ for genes *j* = 1, *…*, *J*. Some of these may be zero, if a gene is not turned on in the cell. Knowing *x*_*ij*_ is equivalent to knowing the total number of transcripts *t*_*i*_ = Σ_*j*_ *x*_*ij*_ and the relative proportions of each gene *π*_*ij*_, since *x*_*ij*_ = *t*_*i*_*π*_*ij*_. The total number of UMI counts *n*_*i*_ = Σ*j y*_*ij*_ does not estimate *t*_*i*_. However, under multinomial sampling, the UMI relative abundances 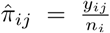 are MLEs for the true proportions *π*_*ij*_. Note that it is possible that 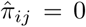 even though *π*_*ij*_ *>* 0, indicating a dropout. Because 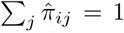 regardless of *n*_*i*_, the use of multinomial MLEs is equivalent to the widespread practice of normalizing each cell by the total counts. Furthermore, the use of size factors *s*_*i*_ = *n*_*i*_*/m* leads to 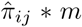 (if *m* = 10^6^ this is CPM).

Traditional bulk RNA-Seq experiments measured gene expression in read counts of many cells per sample rather than UMI counts of single cells. Gene counts from bulk RNA-Seq could thus range over several orders of magnitude. To facilitate comparison of these large numbers many bulk RNA-Seq methods have relied on a logarithm transformation. This enables interpretation of differences in normalized counts as fold changes on a relative scale. Prior to the use of UMIs, scRNA-Seq experiments also produced read counts with wide ranging values, and a log transform was again employed. However, with single cell data, more than 90% of the genes might be observed as exact zeros, and log(0) = – ∞ which is not useful for data analysis. UMI data also contain large numbers of zeros, but do not contain very large counts since PCR duplicates have been removed. Nevertheless, log transformation has been commonly used with UMI data as well.

The current standard is to transform the UMI counts as log_2_ 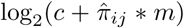 where *c* is a pseudocount to avoid taking the log of zero, and typically *c* = 1. As before, *m* is some constant such as 10^6^ for CPM. Finally, the data are centered and scaled so that the mean of each gene across cells is zero, and the standard deviation is one. This standardization of the data causes any subsequent computation of distances or dimension reduction to be invariant to constant additive or multiplicative scaling. For example, under Manhattan distance *d*(*x* + *c, y* + *c*) = |*x* + *c* − (*y* + *c*) | = |*x − y|* = *d*(*x, y*). In particular, using size factors such as CPM instead of relative abundances leads to a rescaling of the pseudocount, and use of any pseudocount is equivalent to replacing the MLE with the MAP estimator. Let *k* = *c/m* and *α*_*i*_ = *kn*_*i*_. Then the weight term in the MAP formula becomes *w*_*i*_ = *Jk/*(1 + *Jk*) = *w* which is constant across all cells *i*. Furthermore *Jk* = *w/*(1 *– w*), showing that

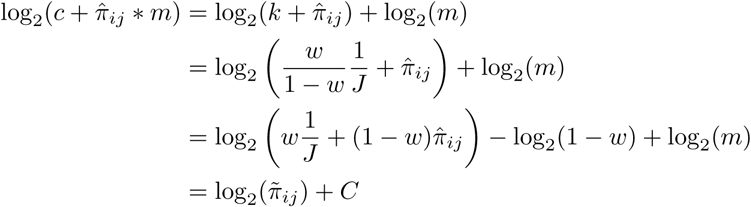

Where C is a global constant that does not vary across cells or genes. For illustration, if *c* = 1 and *m* = 10^6^ this is equivalent to assuming a prior where all genes are equally expressed and for cell *i*, a weight of *w* = *J/*(10^6^ + *J*) is given to the prior relative to the MLE. Since the number of genes *J* is on the order of 10^4^, we have *w ≈.*01. The prior sample size for cell *i* is *Jα*_*i*_ = 10^*–6*^*Jn*_*i*_ *≈.*01* *n*_*i*_ where *n*_*i*_ is the data sample size. The standard transformation is therefore equivalent to using a weak prior to obtain a MAP estimate of the relative abundances, then log-transforming before dimension reduction.

In most scRNA-Seq datasets, the total number of UMIs *n*_*i*_ for some cells *≤ ≤* may be significantly less than the constant *m*. For these cells, the size factors *s*_*i*_ = *n*_*i*_*/m* are less than one. Therefore, after normalization (dividing by size factor), the counts are scaled up to match the target size of *m*. Due to the discreteness of counts, this introduces a bias after log transformation, if the pseudocount is small (or equivalently, if *m* is large). For example, let *c* = 1 and *m* = 10^6^ (CPM). If *n*_*i*_ = 10^4^ for a particular cell, we have *s*_*i*_ = .01. A raw count of *y*_*ij*_ = 1 for this cell is normalized to 1*/.*01 = 100 and transformed to log_2_(1 + 100) = 6.7. For this cell, on the log scale there cannot be any values between zero and 6.7 because fractional UMI counts cannot be observed, and log_2_ (1 + 0) = 0. Small pseudocounts and small size factors combined with log transform arbitrarily exaggerate the difference betwen a zero count and a small nonzero count. As previously shown, this scenario is equivalent to using MAP estimation of *π*_*ij*_ with a weak prior. To combat this distortion, one may attempt to strengthen the prior to regularize 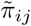 estimation at the cost of additional bias, as advocated by [20]. An extreme case occurs when *c* = 1 and *m* = 1. Here, the prior sample size is *Jn*_*i*_ so almost all the weight is on the prior. The transform is then log_2_.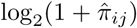 But this function is approximately linear on the domain 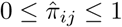. After centering and scaling, a linear transformation is vacuous.

To summarize, log transformation with a weak prior (small size factor, such as CPM) introduces strong artificial distortion between zeros and nonzeros, while log tranformation with a strong prior (large size factor) is roughly equivalent to not log transforming the data.

### 4.3 Generalized PCA

PCA minimizes the mean squared error (MSE) between the data and a lowrank representation, or embedding. Let *y*_*ij*_ be the raw counts and *z*_*ij*_ be the normalized and transformed version of *y*_*ij*_ such as centered and scaled log-CPM (z-scores). The PCA objective function is:

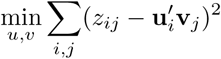

where **u**_*i*_, **v**_*j*_ ∈ ℝ*^L^* for *i* = 1, *…*, *I, j* = 1, *…*, *J*. The **u**_*i*_ are called factors or principal components and the **v**_*j*_ are called loadings. The number of latent dimensions *L* controls the complexity of the model. Minimization of the MSE is equivalent to minimizing the Euclidean distance metric between the embedding and the data. It is also equivalent to maximizing the likelihood of a Gaussian model

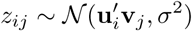

If we replace the Gaussian model with a Poisson, which approximates the multinomial, we can directly model the UMI counts as

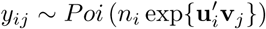

or alternatively, in the case of overdispersion, we may approximate the Dirichlet-multinomial using a negative binomial likelihood

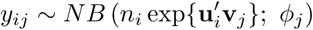

We define the *linear predictor* as 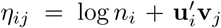. It is clear that the mean 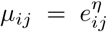 appears in both the Poisson and Negative Binomial model statements, showing that the latent factors interact with the data only through the mean. We may then estimate **u**_*i*_, **v**_*j*_ (and *φ*_*j*_) by maximizing the likelihood (in practice, adding a small L2 penalty to large parameter values improves numerical stability). A link function must be used since **u**_*i*_, **v**_*j*_ are real valued whereas the mean of a Poisson or negative binomial must be positive. The total UMIs *n*_*i*_ term is used as an offset since no normalization has taken place; alternative size factors *s*_*i*_ such as those from scran [19] could be used in place of *n*_*i*_. If the first element of each **u**_*i*_ is constrained to equal 1, this induces a gene-specific intercept term in the first position of each **v**_*j*_, which is analogous to centering. Otherwise, the model is very similar to that of PCA; it is simply optimizing a different objective function. Unfortunately, MLEs for **u**_*i*_, **v**_*j*_ cannot be expressed in closed form, so an iterative Fisher Scoring procedure is necessary. We refer to this model as GLM-PCA. Just as PCA minimizes MSE, GLM-PCA minimizes a generalization of MSE called the *deviance* [48]. While generalized PCA has been discovered before by [29], our implementation is novel in that it allows for intercept terms, offsets, and non-canonical link functions. We also use a blockwise update for optimization which we found to be more numerically stable than that of [29]; we iterate over latent dimensions *l* rather than rows or columns. This technique is inspired by non-negative matrix factorization algorithms such as hierarchical alternating least squares and rank-one residue iteration, see [49] for a review.

As an illustration, consider GLM-PCA with the Poisson approximation to a multinomial likelihood. The objective function to be minimized is simply the overall deviance:

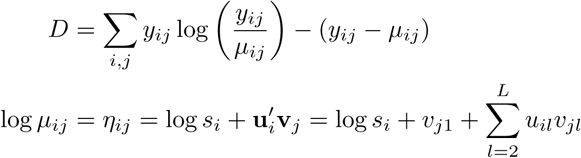

where *s*_*i*_ is a fixed size factor such as the total number of UMIs (*n*_*i*_). The optimization proceeds by taking derivatives with respect to the unknown parameters: *v*_*j*1_ is a gene-specific intercept term, and the remaining *u*_*il*_, *v*_*jl*_ are the latent factors.

The GLM-PCA method is most concordant to the data generating mechanism since all aspects of the pipeline are integrated into a coherent model rather than being dealt with through sequential normalizations and transformations. The interpretation of the **u**_*i*_ and **v**_*j*_ vectors is the same as in PCA. For example, suppose we set the number of latent dimensions to two (i.e. *L* = 3 to account for the intercept). We can plot *u*_*i*2_ on the horizontal axis and *u*_*i*3_ on the vertical axis for each cell *i* to visualize relationships between cells such as gradients or clusters. In this way, the **u**_*i*_ and **v**_*j*_ capture biological variability such as differentially expressed genes.

### 4.4 Residuals and z-scores

Just as mean squared error can be computed by taking the sum of squared residuals under a Gaussian likelihood, the deviance is equal to the sum of squared *deviance residuals* [48]. Since deviance residuals are not well-defined for the multinomial distribution, we adopt the binomial approximation. The deviance residual for gene *j* in cell *i* is given by

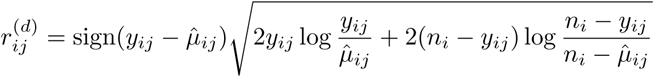

where under the null model of constant gene expression across cells, 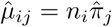. The deviance residuals are the result of regressing away this null model. An alternative to deviance residuals is the Pearson residual, which is simply the difference in observed and expected values scaled by an estimate of the standard deviation. For the binomial, this is

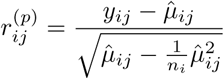

According to the theory of generalized linear models (GLM), both types of residuals follow approximately a normal distribution with mean zero if the null model is correct [48]. Deviance residuals tend to be more symmetric than Pearson residuals. In practice, the residuals may not have mean exactly equal to zero, and may be standardized by scaling their gene-specific standard deviation just as in the Gaussian case.

The z-score is simply the Pearson residual where we replace the multinomial likelihood with a Gaussian (normal) likelihood, and use normalized values instead of raw UMI counts. Let *q*_*ij*_ be the normalized (possibly log-transformed) expression of gene *j* in cell *i* without centering and scaling. The null model is that the expression of the gene is constant across all cells:

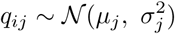

The MLEs are 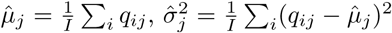, and the z-scores equal the Pearson residuals 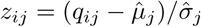.

### 4.5 Feature selection using deviance

Genes with constant expression across cells are not informative. Such genes may be described by the multinomial null model where *π*_*ij*_ = *π*_*j*_. Goodness of fit to a multinomial distribution can be quantified using deviance, which is twice the difference in log-likelihoods comparing a saturated model to a fitted model. The multinomial deviance is a joint deviance across all genes and for this reason is not helpful for screening informative genes. Instead, one may use the binomial deviance as an approximation:

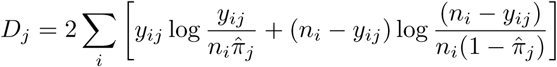

A large deviance value indicates the model in question provides a poor fit. Those genes with biological variation across cells will be poorly fit by the null model and will have the largest deviances. By ranking genes according to their deviances, one may thus obtain highly deviant genes as an alternative to highly variable or highly expressed genes.

### 4.6 Systematic Comparison of Methods

We considered combinations of the following methods and parameter settings, following [14]. Italics indicate methods proposed in this manuscript. Feature selection: highly expressed genes, highly variable genes, and *highly deviant genes*. We did not compare against highly dropout genes because [14] found this method to have poor downstream clustering performance for UMI counts and it is not as widely used in the literature. Number of genes: 60, 300, 1,500. Normalization, transformation, and dimension reduction: PCA on log-CPM zscores, ZINB-WAVE [27], *PCA on deviance residuals, PCA on Pearson residuals*, and *GLM-PCA*. Number of latent dimensions: 10, 30. Clustering algorithm: k-means [50], Seurat [16]. Number of clusters: all values from 2-10, inclusive. Seurat resolution: 0.05, 0.1, 0.2, 0.5, 0.8, 1, 1.2, 1.5, 2.

## 5. Declarations

### Ethics approval and consent to participate

Not applicable.

### Consent for publication

Not applicable.

### Availability of data and material

All methods and assessments described in this manuscript are publicly available at https://github.com/willtownes/scrna2019.

### Authors’ contributions

SCH, MJA, and RAI identified the problem. FWT proposed, derived, and implemented the GLM-PCA model, its fast approximation using residuals, and feature selection using deviance. SCH, MJA and RAI provided guidance on refining the methods and evaluation strategies. FWT and RAI wrote the draft manuscript and revisions were suggested by SCH and MJA. All authors approved the final manuscript.

## Acknowledgements

The authors thank Keegan Korthauer, Jeff Miller, Linglin Huang, Alejandro Reyes, and Yered Pita-Juarez for valuable suggestions.

## Supplemental Figures

**Figure S1:**
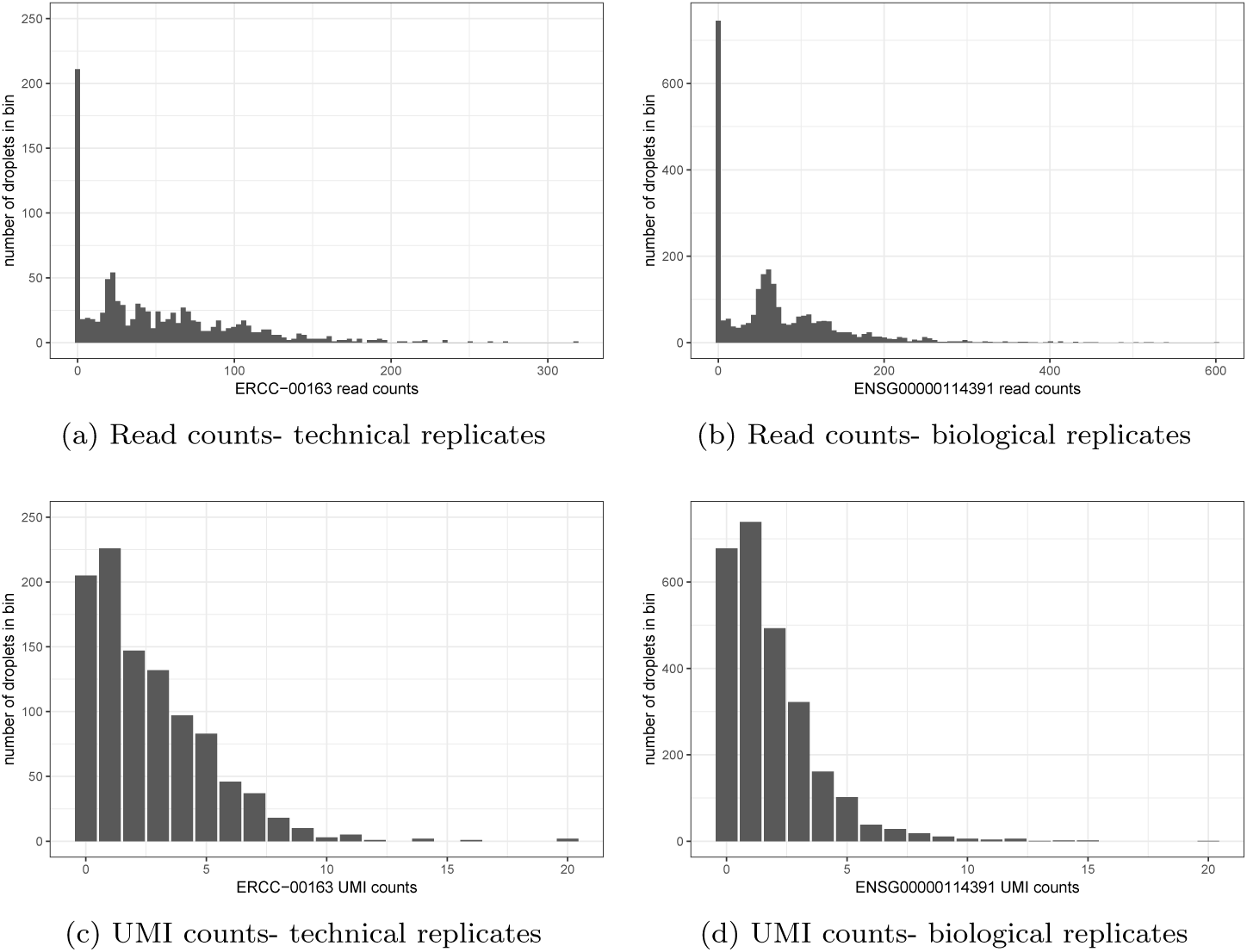
Comparing read counts and UMI counts sampling distribution from technical and biological replicates negative control datasets. a) Read count distribution for spike-in ERCC-00163 across technical replicates. b) Read count distribution for gene ENSG00000114391 across biological replicates (purified monocytes). c) as a) but without PCR duplicates. d) as b) but without PCR duplicates.

**Figure S2:**
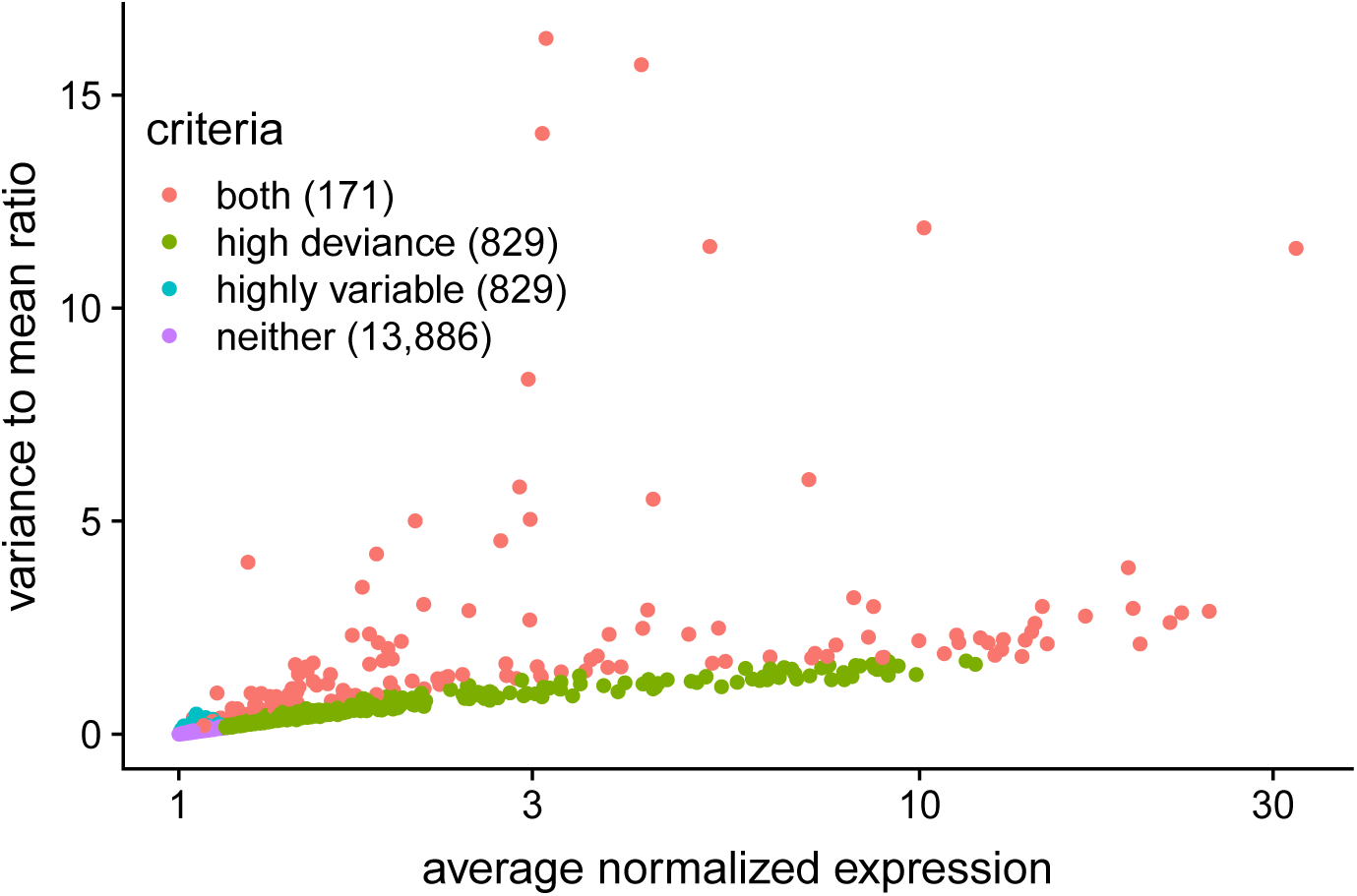
Comparison of top 1,000 genes selected as most informative in the Zheng 8eq dataset. The variance to mean ratio for each gene is plotted against the average expression. Counts were normalized using scran [19]. Colors represent genes that are in the top 1,000 ranked by variability (blue, red) and top 1,000 ranked by approximate multinomial deviance (green, red). Red indicates genes identified by both criteria, while purple indicates genes identified by neither criteria. Note that highly expressed genes have large values on the horizontal axis. The number of genes in each category is shown in parentheses.

**Figure S3:**
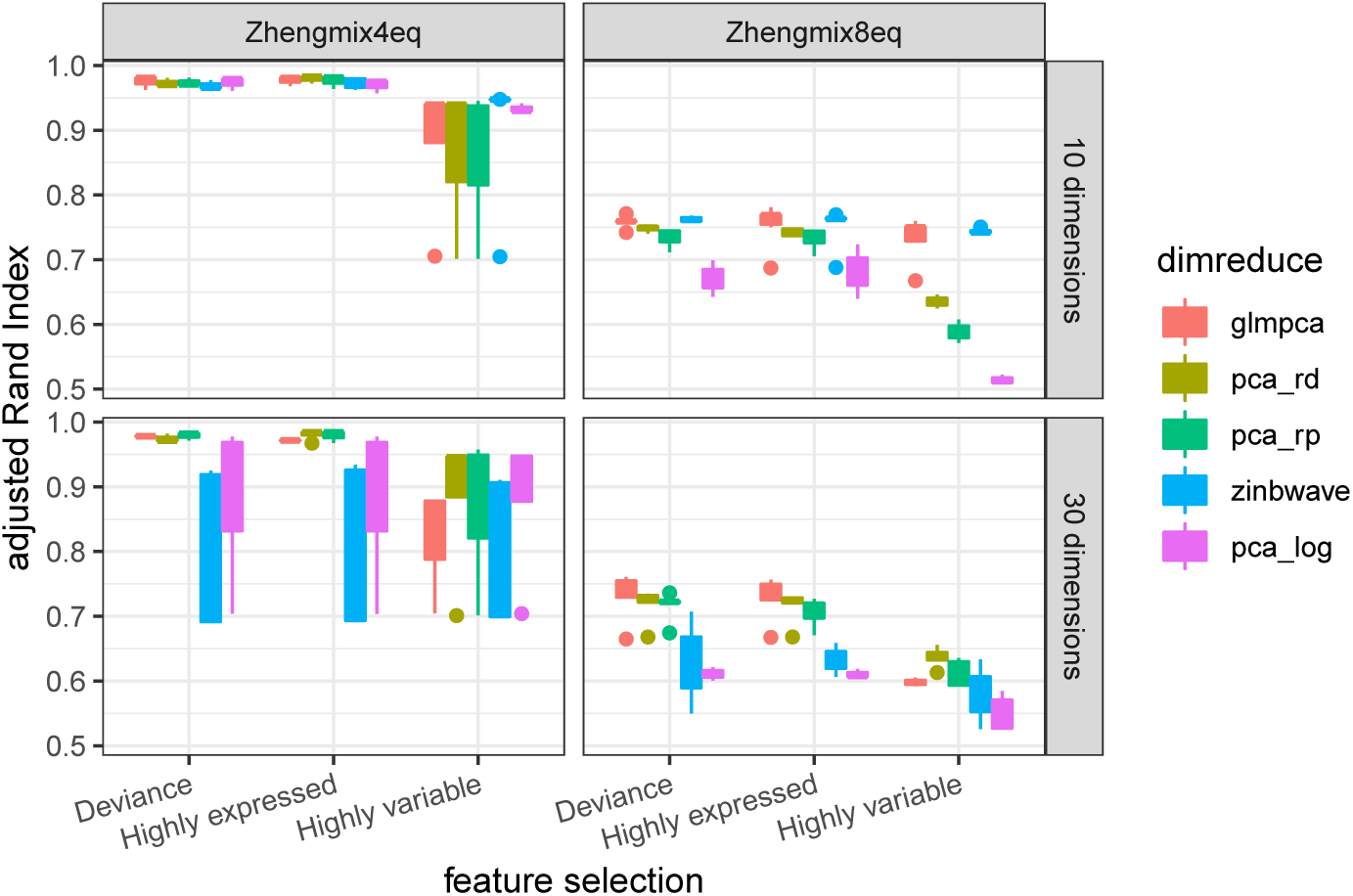
Comparison of Seurat clustering performance for all dimension reduction and feature selection methods on ground-truth datasets from [14]. The number of informative genes was fixed at 1,500. The Poisson approximation to the multinomial was used for GLM-PCA. Only results with the number of clusters within 25% of the true number are presented. Abbreviations: dimreduce: dimension reduction method, pca rd: PCA on deviance residuals, pca rp: PCA on Pearson residuals, pca log: PCA on log-CPM.

**Figure S4:**
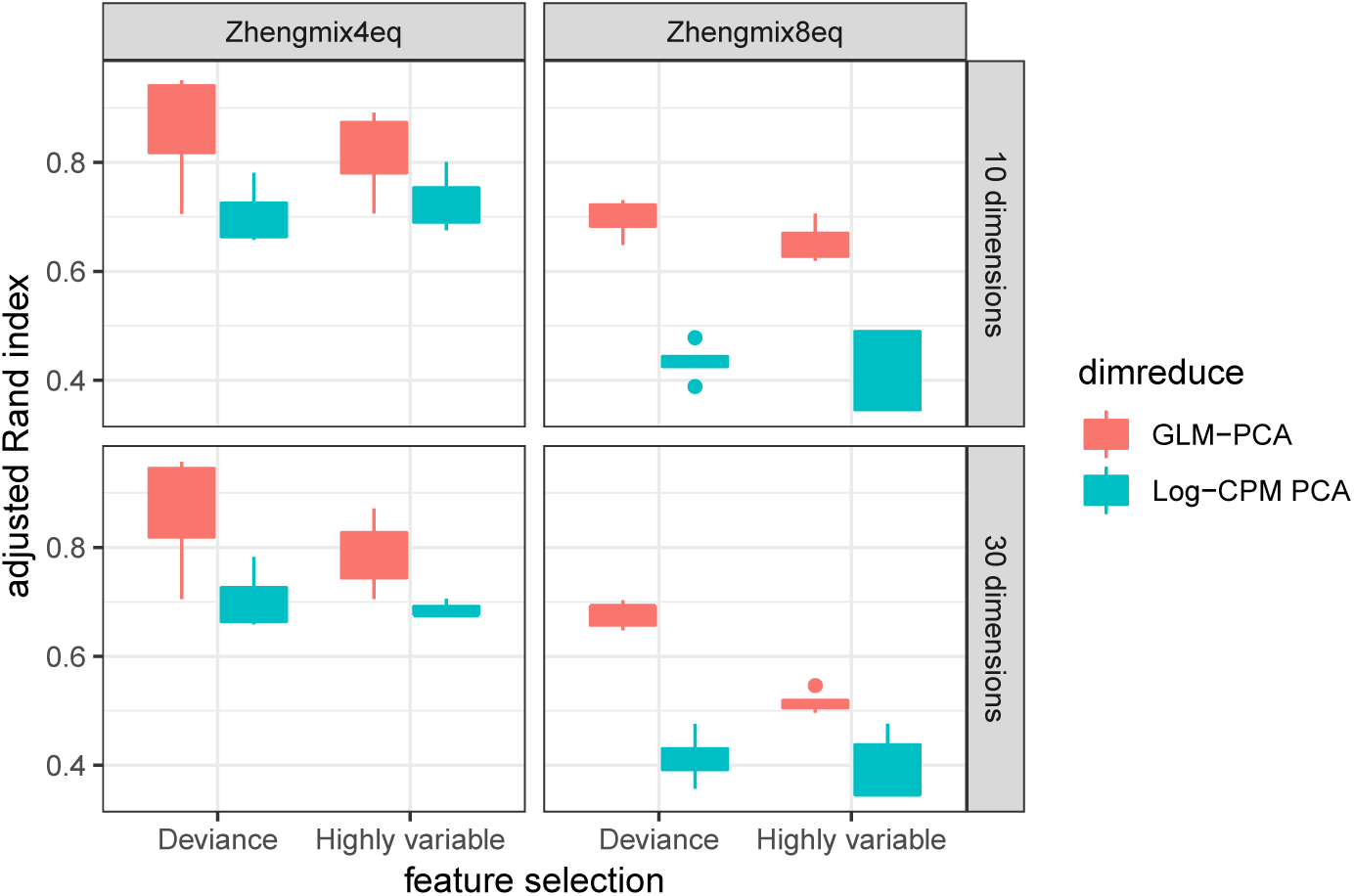
Dimension reduction with GLM-PCA and feature selection using deviance improves k-means clustering performance. Each column represents a different ground-truth dataset from [14]. The top 1,500 informative genes were identified by approximate multinomial deviance and highly variable genes. The Poisson approximation to the multinomial was used for GLM-PCA. Only results with the number of clusters within 25% of the true number are presented.

**Figure S5:**
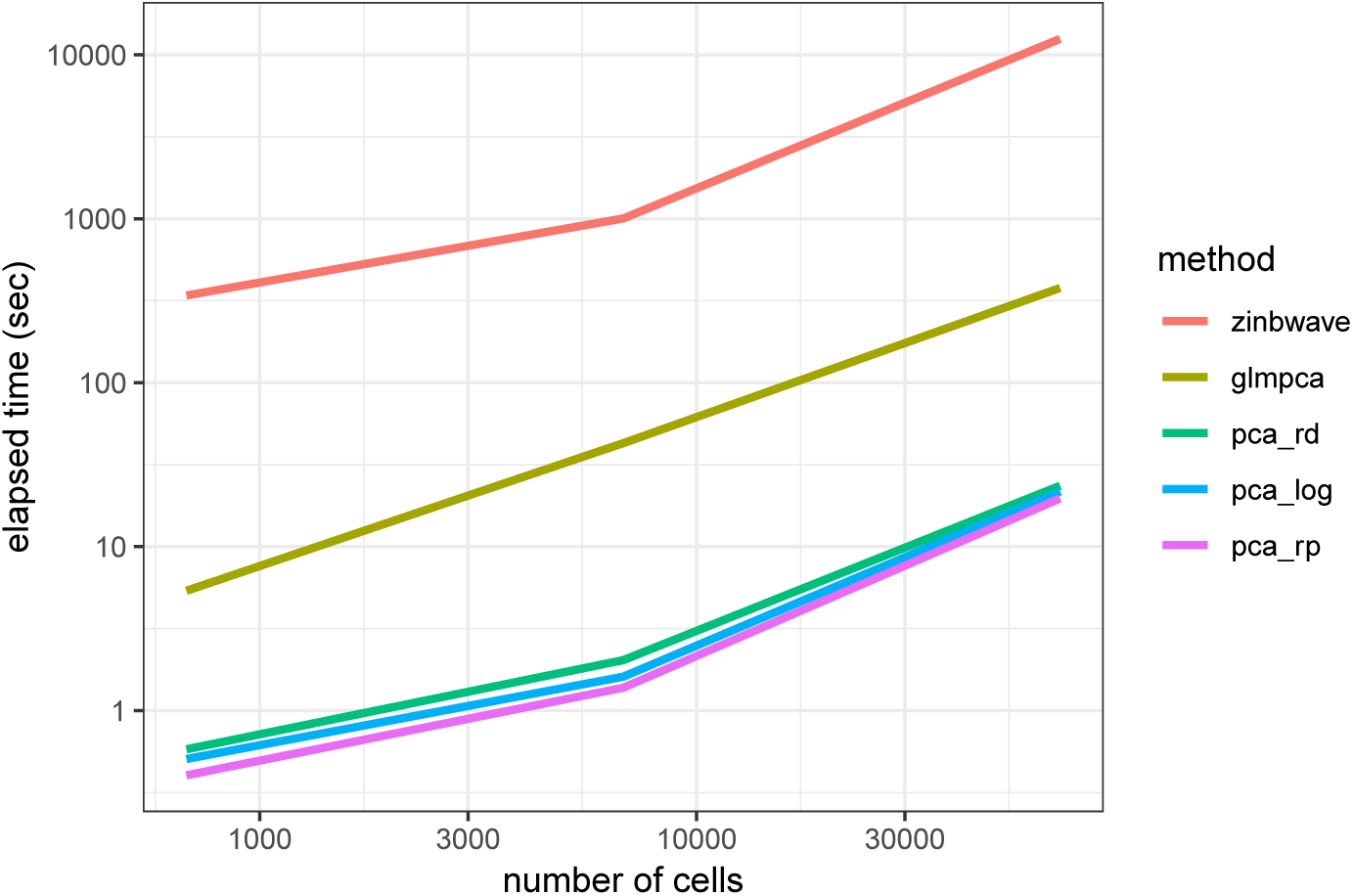
Computational speed comparison of dimension reduction methods GLM-PCA (glmpca), ZINB-WAVE (zinbwave), PCA on deviance residuals (pca rd), PCA on Pearson residuals (pca rp), and PCA on log-CPM (pca log).

